# Brain Morphometry and Chronic Inflammation in Bangladeshi Children Growing up in Extreme Poverty

**DOI:** 10.1101/2021.12.26.474220

**Authors:** Ted K. Turesky, Laura Pirazzoli, Talat Shama, Shahria Hafiz Kakon, Rashidul Haque, Nazrul Islam, Amala Someshwar, Borjan Gagoski, William A. Petri, Charles A. Nelson, Nadine Gaab

## Abstract

Over three hundred million children live in environments of extreme poverty, and the biological and psychosocial hazards endemic to these environments often expose these children to infection, disease, and inflammatory responses. Chronic inflammation in early childhood has been associated with diminished cognitive outcomes, and despite this established relationship, the mechanisms explaining how inflammation affects brain development are not well known. Importantly, the prevalence of chronic inflammation in areas of extreme poverty raises the possibility that it may serve as a mechanism explaining the known relationship between low socioeconomic status (SES) and altered brain development. To examine these potential pathways, seventy-nine children growing up in an extremely poor, urban area of Bangladesh underwent MRI scanning at six years of age. Structural brain images were submitted to Mindboggle software, a Docker-compliant and high-reproducibility tool for regional estimations of volume, surface area, cortical thickness, sulcal depth, and mean curvature. C-reactive protein was assayed at eight time points between infancy and five years of age and the frequency with which children had elevated concentrations of inflammatory marker represented the measure of chronic inflammation. Childhood SES was measured with maternal education and income-to-needs (i.e., monthly household income divided by the number of household members). Chronic inflammation predicted volume in bilateral basal ganglia structures and mediated the link between maternal education and bilateral putamen volumes. These findings suggest that chronic inflammation is associated with brain morphometry in the basal ganglia, predominantly the putamen, and further offers inflammation as a potential mechanism linking SES to brain development.

## INTRODUCTION

Roughly one in six children worldwide grow up in extreme poverty (UNICEF; https://www.unicef.org/social-policy/child-poverty), where biological and psychosocial hazards put them at heightened risk for infection, disease, and consequent inflammatory responses. Inflammation early in life can have a profound impact on neurodevelopment and has been associated with diminished cognitive outcomes (Jiang et al., 2018). For instance, higher concentrations of inflammatory markers in 6-month-old infants have been associated with subsequent lower cognitive, language, motor, and socio-emotional outcomes at age two years (Jiang et al., 2014, 2017). Chronic or sustained inflammation, measured as the number or frequency of times a child has elevated concentrations of inflammatory markers, has also been examined in the context of neurodevelopment (Jiang et al., 2017; Donowitz et al., 2018). Rather than focus on specific inflammatory markers, this approach has instead employed examining the inflammatory marker C-reactive protein (CRP): more specifically, the frequency of CRP elevations in the first two years of life was associated with lower neurocognitive outcomes at age two years (Jiang et al., 2017) and mediated the relationship between poverty and verbal and performance IQ at age five years (Jensen et al., 2019a).

This relationship between inflammation and neurocognitive outcomes has strong mechanistic support in studies examining the effects of inflammation on brain anatomy and physiology in non-human animal models. This body of evidence has shown that, first, inflammatory markers need to penetrate the central nervous system (CNS), which occurs through increased permeability of the blood-brain barrier (BBB) during conditions of systemic inflammation (Sharshar et al., 2005; Diamond et al., 2013; Sankowski et al., 2015; Varatharaj and Galea, 2017). Evidence suggests that circumventricular organs and dural venous sinuses could also serve as fenestrations for inflammatory markers into the CNS (Popescu et al., 2011; Fitzpatrick et al., 2020). The translocation of inflammatory markers into the CNS can then alter the action of glial cells, which ordinarily serve as mediators of neuroinflammation. In particular, microglia, as the macrophages of the brain, are responsible for local responses to injury and infection. Inflammatory markers have been shown to activate microglia (Sankowski et al., 2015) and potentially in a region-specific manner; e.g., greater numbers of activated microglia were detected in the putamen, hippocampus, and cerebellum in post-mortem tissue from individuals with sepsis compared with controls (Westhoff et al., 2019). This has important implications as activated microglia can trigger neuronal death (Bessis et al., 2007), which may in turn reduce gray and white matter tissue. Bolstering this, activation of microglia by administering inflammatory markers has been shown to selectively degenerate dopaminergic neurons in the basal ganglia (Gao et al., 2002). Additionally, during sustained inflammation, astrocytes have been shown to upregulate inflammatory processes and inhibit neuronal repair and axonal growth (Jerry Silver et al., 2006). Lastly, early exposure to systemic inflammatory cytokine has been shown to disrupt oligodendrocyte maturation, reducing white matter myelination and consequently estimates of fractional anisotropy (Favrais et al., 2011). Taken together, inflammation may alter brain structure and function by increasing translocating into the CNS and altering glia interactions.

Inflammation has also been linked with brain structure and function in humans. For example, white matter damage was more common in infants with elevated concentrations of inflammatory markers in umbilical blood (Duggan et al., 2001; Hansen-Pupp et al., 2005) and amniotic fluid (Yoon et al., 1997). In addition, higher maternal inflammatory marker concentration during pregnancy related to greater volume in the right amygdala (Graham et al., 2018) as well as alterations in functional connectivity in multiple brain networks/regions in infants (Rudolph et al., 2018), and even in adults 45 years later (Goldstein et al., 2021). In adults, greater concentrations of inflammatory markers were cross-sectionally associated with less total intracranial volume (Jefferson et al., 2007), less hippocampal volume (Marsland et al., 2008), less mean white matter fractional anisotropy (Gianaros et al., 2013), and weaker functional connectivity in neural networks associated with emotion regulation and executive functioning (Nusslock et al., 2019).

Structural and functional brain alterations have also been observed in diseases and conditions related to inflammation. For example, multiple sclerosis, a neurodegenerative and inflammatory disorder (Compston and Coles, 2008), is predominantly characterized by white matter demyelination, but also by gray matter atrophy (Bø et al., 2003; Tiberio et al., 2005; Vercellino et al., 2005; Fisher et al., 2008) and reduced volume mainly in left hemisphere cortical regions and bilateral caudate (Prinster et al., 2006). Also, inflammatory marker concentration has been associated with white matter fractional anisotropy in adults with major depressive disorder (Sugimoto et al., 2018). In adults hospitalized with sepsis, which is characterized by a heightened systemic inflammatory response (Bosmann and Ward, 2013), less cerebral gray and white matter, and deep gray matter volume has been observed compared with controls (Orhun et al., 2018, 2019). In adults with hepatitis C virus infection and malignant melanomas, administration of inflammatory marker has been associated with altered dopamine function, glucose metabolism, and activation assayed with functional MRI in the basal ganglia (Capuron et al., 2007, 2012). While it is not entirely clear why this inflammation exhibits spatial specificity for the basal ganglia, there seems to be abundant evidence from a wide range of human and non-human animal experiments suggesting that dopaminergic systems are particularly susceptible to inflammatory activity (Shuto et al., 1997; Kamata et al., 2000; Capuron et al., 2007, 2012), which agrees with the hypothesis that inflammation affects cortico-basal ganglia circuits (Nusslock and Miller, 2016).

To a lesser extent, the relationship between inflammation and the brain has also been studied in the context of extreme poverty. This is an important field of inquiry to bolster, as hundreds of millions of children live in impoverished circumstances that may predispose them to inflammation early in life (Bhutta et al., 2017; John et al., 2017; Kutlesic et al., 2017; Nelson, 2017; Rondó et al., 2013). Here, children are exposed to water and air pollution, as well as organic and inorganic waste. They also experience food scarcity and malnutrition, which when coupled with illness, prolong elevated inflammatory states (Rytter et al., 2014); e.g., children experience immune dysfunction when bereft of key nutrients. In turn, elevated inflammatory states are associated with worsening malnutrition as enteric diseases, which are prevalent in low-resource settings and accompanied by inflammation, often reduce appetite (Jensen et al., 2017). Additionally, a large subset of these children experience myriad psychosocial stressors (e.g., family conflict; Nelson 2017), which can also affect inflammatory processes (Chiang et al., 2015). Taken altogether, these biological and psychosocial hazards predispose children from low-resource settings to conditions of chronic inflammation.

Neuroimaging studies of *chronic*, as opposed to acute, inflammation are rare because they require longitudinal data collection over many years and because chronic inflammation is most prevalent in parts of the world that often do not have access to brain imaging equipment for research purposes. Consequently, only two studies examined chronic inflammation in the context of brain development, both utilizing electroencephalography (EEG) and both conducted in Bangladesh, where unhygienic conditions imperil child health and safety in a manner that is extremely hazardous compared with high-resource settings (Nelson, 2017). They found that the frequency of CRP elevations during the first two years of life predicted brain function at three years of age, particularly, neural responses to faces (Xie et al., 2019a) and parietal-occipital functional connectivity (Bach et al., 2022). To summarize, inflammation seems to be associated with alterations in brain structure and function in widespread brain areas, in gray and white matter, and with various metrics (e.g., volume and presence/absence of damage).

Further, as chronic inflammation is of major concern in low-resource settings (Bhutta et al., 2017; John et al., 2017; Kutlesic et al., 2017; Nelson, 2017), it is also important to consider the role that socioeconomic status (SES) might play in the relationship between inflammation and brain development. As with inflammation, SES in general has also been linked to brain structure (Noble et al., 2012; Hanson et al., 2013; Lawson et al., 2013; Luby et al., 2013; Hair et al., 2015; Mackey et al., 2015; Noble et al., 2015; McDermott et al., 2019; Spann et al., 2020). However, such links are inherently distal (Farah, 2017, 2018), as SES does not affect the brain directly, but instead influences a wide array of biological and psychosocial risk and protective factors, each of which could affect brain development independently or in combination (Jensen et al., 2017). Inflammation is one of the more frequently examined biological factors in the context of brain development (Yoon et al., 1997; Duggan et al., 2001; Hansen-Pupp et al., 2005; Graham et al., 2018; Rudolph et al., 2018; Goldstein et al., 2021), but others, such as malnutrition (Gunston et al., 1992; Hazin et al., 2007; Atalabi et al., 2010; El-Sherif et al., 2012; Kumar et al., 2016; Lelijveld et al., 2019) and air pollution (Calderón-Garcidueñas et al., 2011), have also been investigated. In contrast, psychosocial risk factors, including parental mental health difficulties and deprivation, have been recurrently associated with brain development (Rao et al., 2010; Tottenham et al., 2010; Sheridan et al., 2012; McLaughlin et al., 2014; Lebel et al., 2016; Wen et al., 2017; Merz et al., 2020; VanTieghem et al., 2021) and methodically conceptualized (Sheridan and McLaughlin, 2014; Tooley et al., 2021). Overall, many of the risk factors associated with SES have been linked with brain development, and it is possible that a subset of these *could* serve as mechanistic intermediaries in observed links between SES and brain development.

However, there remains a paucity of formal mediation testing to determine which risk factors mediate the link between SES and brain development. This is an important gap to address, as mediation testing is a necessary (though not entirely sufficient) precursor to establish causal pathways (Farah, 2017). Among these few studies that have conducted formal mediation testing, stress and caregiving quality have been shown to mediate the relation between SES and hippocampal volume (Luby et al., 2013). Similarly, cortisol, the canonical stress hormone, has been shown to mediate the relation between SES and CA3 and dentate gyrus (hippocampus) volumes (Merz et al., 2019). Also, the home linguistic environment was found to mediate the association between SES and left perisylvian cortical surface area (Merz et al., 2020). With regard to biological hazards, diminished growth, a proxy for malnutrition, mediates the link between SES and volume mainly in subcortical gray matter and white matter regions (Turesky et al., 2021). CRP concentration has also been shown to mediate the link between white matter organization as measured with diffusion imaging, and smoking and adiposity, two hazards themselves linked with SES, but CRP did not mediate the link between SES and white matter organization directly (Gianaros et al., 2013). However, CRP concentration here was measured at a single time point in adulthood and given the mediating role of chronic CRP elevation between poverty and neurocognitive outcomes (Jensen et al., 2019a), it is more likely that repeated measurements of CRP across development would, in contrast, represent a mediator to the link between poverty and brain development. Ultimately, identifying mediating risk factors is an important first step for preventing or remedying adverse effects of SES disadvantage (Olson et al., 2021).

The goals of the present study were two-fold. First, we examined whether chronic inflammation was related to global and regional measures of brain morphometry in Bangladeshi children, a population largely unrepresented in developmental cognitive neuroscience. Chronic inflammation, collected at eight time points between birth and age five years, was approximated as the frequency with which children’s CRP levels were elevated during this important time of brain development; this method for estimating chronic inflammation is well-established (Naylor et al., 2015; Jiang et al., 2017; Jensen et al., 2019b; Xie et al., 2019b; Bach et al., 2022). Measures of volume, surface area, cortical thickness, sulcal depth, and mean curvature were estimated by submitting structural MRI images acquired at six years of age to Mindboggle (Klein et al., 2017), using a Docker container for reproducibility. Second, we examined whether chronic inflammation mediated the links between SES and brain morphometry that were observed in a previous study (Turesky et al., 2021). Based on past work linking brain volume to inflammation or inflammatory conditions (Jefferson et al., 2007; Orhun et al., 2018, 2019) and the links that SES shares with chronic inflammation (Jensen et al., 2019b) and brain volume (Noble et al., 2012; Hanson et al., 2013; Luby et al., 2013; Hair et al., 2015; Noble et al., 2015; McDermott et al., 2019), we hypothesized that chronic inflammation would correlate with brain volume and also mediate links between SES and brain volume. As with earlier neuroscientific reports, we further hypothesized that these associations would favor subcortical gray matter regions, such as basal ganglia (e.g., Capuron et al., 2012, 2007) and medial temporal lobe structures (Graham et al., 2016) and white matter (Duggan et al., 2001; Hansen-Pupp et al., 2005; Orhun et al., 2018, 2019), as these regions have been associated with inflammation or inflammatory conditions.

The extant literature does not motivate hypotheses for associations between chronic inflammation and surface-based measures. However, given associations between inflammatory conditions and several cortical areas (Prinster et al., 2006; Orhun et al., 2018, 2019), and the sensitivity of surface-based measures to SES (Lawson et al., 2013; Mackey et al., 2015; Noble et al., 2015; McDermott et al., 2019) and early developmental injuries (Shimony et al., 2016), it is possible that these measures would also relate to chronic inflammation. Therefore, we also examined the relation between chronic inflammation and global and regional surface-based estimates in an exploratory manner.

## METHODS

### Participants

The work presented here is a subset of the Bangladesh Early Adversity Neuroimaging study (Donowitz et al., 2018; Jensen et al., 2019a, 2019b; Moreau et al., 2019; Turesky et al., 2019, 2020, 2021; Xie et al., 2019a, 2019b; Bach et al., 2022), which examines early brain development in children exposed to extreme poverty in Dhaka, Bangladesh. Beginning in infancy, neuroimaging, socioeconomic, anthropometric, behavioral, and biological measures were collected in 130 children. Of these children, 81 underwent structural MRI between five and seven years of age. Severe motion artifacts were present in two participants’ datasets, thereby decreasing the final sample to 79 structural MRI datasets (6.68 ± 0.40 years; F/M = 36/43). All datasets were evaluated by a clinical radiologist in Bangladesh and a pediatric neuroradiologist in the United States to ensure the absence of malignant brain features. Additionally, no child had been diagnosed with a neurological disorder or disease. The study was approved by research and ethics review committees at the institutions affiliated with the authors.

### Chronic Inflammation

Inflammation was initially measured as the concentration of serum C-reactive protein (CRP) in mg/L. Children in this study had CRP assayed at eight time points over the first five years of life:at 6, 18, 40, 53, 104, 156, 207 and 260 weeks. Inflammation was initially measured as the concentration of serum C-reactive protein (CRP) in mg/L. Children in this study had CRP assayed at eight time points over the first five years of life: at 6, 18, 40, 53, 104, 156, 207 and 260 weeks; only children with CRP estimates at all eight time points were included in the final sample. Descriptive statistics and histograms for CRP at each time point are provided in Supplementary Table 1 and Supplementary Figure 1, respectively. Overall, median values reported here are considerably higher than 50^th^ percentile values reported in previous literature for U.S. (Ford et al., 2003) and European children (Schlenz et al., 2014), which are roughly 0.3 mg/L.

In line with previous work with this cohort, chronic inflammation was measured in each child as the number of times their CRP concentration was elevated across the number of measurements (Jensen et al., 2019b; Xie et al., 2019b; Bach et al., 2022). At each measurement, a child’s CRP concentration was designated as elevated when it was greater than the group median (Naylor et al., 2015; Jiang et al., 2017; Xie et al., 2019a). As such, the CRP index ranges between 0 and 8, with higher values indicating greater chronic inflammation. It is also important to note that for consistency with other studies conducted by our lab with other neuroimaging methods, the median value was based on a larger cohort of 130 children. The children in the current study, who represent a subset of the larger cohort, had on average 4 ± 2 CRP elevations.

### Measures of socioeconomic status (SES)

Maternal education and income-to-needs, calculated from years of mother’s formal education, monthly family income, and number of household members, were used as measures of SES. Whereas maternal education is more strongly related to parenting, which includes cognitive and socioemotional stimulation conferred by the home learning environment (Hoff et al., 2002), income-to-needs is thought to reflect resource availability (Brito and Noble, 2014). Treating these measures as range, graded variables is consistent with current approaches in neuroimaging studies (Lawson et al., 2013; Noble et al., 2015; Betancourt et al., 2016; Brito et al., 2016; Merz et al., 2018) and recommendations for examining SES (Adler et al., 1994; Duncan et al., 2017). Maternal education was estimated along an ordinal scale, ranging from 0 to 10, in which 0 indicates no formal education, 1–9 indicate number of grades passed, and 10 indicates education beyond the 9th grade. On average, mothers of infants in the present study had 4.4 ± 3.9 years of education. Income-to-needs was calculated as the monthly family income divided by the number of household members. On average, these households had an income-to-needs ratio of 3100 ± 1700 Tk (Taka; Bangladeshi currency). Importantly, this is below the World Bank international standard for extreme poverty of Tk4800 per month per household measure, as computed using the USD$1.90 per day per person standard (https://data.worldbank.org/) and an exchange rate of USD$1:Tk85. Individually, 71 children were growing up in extreme poverty. Each SES variable was measured at 6 months and at 3 years of age by interviewing the children’s parents and no children were missing data for either variable at either time point. Measures were then averaged across time points to better capture the measures of SES across childhood. Measures acquired at 6 and 36 months of age were strongly correlated for maternal education (r = 0.94, p = 2.4 x 10^-37^) and income-to-needs (r = 0.67, p = 8.7 x 10^-12^). Due to a positive skew in the income-to-needs variable, these data were log (base 10) transformed prior to statistical analyses.

### Anthropometry

Height-for-age (HAZ) scores were computed from height (in centimeters), as measured by trained, local staff, age (in years), and biological sex. Height measures were submitted to the Anthro Plus software (https://www.who.int/growthref/tools/en/; Multicenter Growth Reference Study; de Onis et al., 2004) and compared with standard growth curves derived from 8440 infants (0–24 months) and children (18–71 months) from Brazil, Ghana, India, Norway, Oman and the U.S. This produced standardized estimates that quantified age- and sex-referenced height deviations from the typical growth trajectory. Importantly, the infants and children used to compute the standard growth curves were raised in environments without severe hazards (e.g., infants were breastfed and not exposed to smoke). Such environments are “likely to favour the achievement of their full genetic growth potential” (de Onis et al., 2006). Accordingly, deviations from standard growth curves can imply the presence of environmental hazards during upbringing. HAZ estimates from 21, 30, and 36 months were averaged across developmental stages to ensure stability of these measures, though there was already high consistency among these estimates as indicated by intercorrelations among the three combinations of pairwise comparisons (r_avg_ = 0.93, p_avg_ = 2.8 x 10^-32^). On average, children in the present study were slightly above the HAZ = −2 threshold for stunting (Grantham-McGregor et al., 2007); however, when examined individually, 24 children qualified as stunted.

### MRI data acquisition

The MRI dataset used in the present study is one from a previous publication by our group (Turesky et al., 2021) and is publicly available at https://openneuro.org/datasets/ds003877/versions/1.0.1. All neuroimaging data were collected on a 3T Siemens MAGNETOM Verio scanner using a 12-channel head coil at the National Institute for Neuroscience in Dhaka, Bangladesh. Structural T1-weighted magnetization-prepared rapid gradient-echo (MPRAGE) scans were acquired with the following parameters: Repetition time = 2500 ms, echo time = 3.47 ms, 176 sagittal slices, 1 mm^3^ voxels, field of view = 256 mm. Functional and diffusion scans were also acquired but are outside of the scope of this report. Consent forms were completed by parents on the day prior to the scan and head circumference was measured on the day of the scan.

There were additional considerations for scanning in Dhaka. Local staff, who had not previously participated in a large-scale pediatric neuroimaging study involving MRI scans, traveled to the United States for training. In Dhaka, staff escorted participants between their homes and the scanning facility. Upon arriving at the scanning facility, children were taught about the MRI machine and the images it can produce. They then practiced remaining motionless inside and outside a cardboard mock scanner. For further details about the scanning procedure that are unique to pediatric MRI studies in Dhaka, please see Turesky et al., (2021).

### MRI data processing

Data used here were originally processed for our previous study (Turesky et al., 2021), but we briefly describe the processing steps taken here. Images were visually inspected for artifacts. Following the removal of two artifactual datasets, the remaining raw MPRAGE images were submitted to Mindboggle 1.3.8 and run in a Docker container (Klein et al., 2017; https://Mindboggle.readthedocs.io/en/latest/). This pipeline implements Advanced Normalization Tools (ANTs) and FreeSurfer (v6.0.0). First, Mindboggle calls antsCorticalThickness, which includes brain extraction, N4 bias correction, tissue segmentation, and cortical thickness estimation. Subsequently, Mindboggle submits raw images to FreeSurfer’s recon-all, which segments the brain into different tissue classes, approximates pial surfaces, and labels volumes and surfaces according to the Aseg (Fischl et al., 2002) and Desikan-Killiany-Tourville (Klein and Tourville, 2012) atlases for subcortical and cortical regions, respectively. Next, segmentations from ANTs and FreeSurfer are hybridized to rectify tissue mislabeling common to each tool. Outputs from Mindboggle include volumetric measures computed for cortical and subcortical brain regions (including white matter) and surface-based measures such as surface area, cortical thickness (from FreeSurfer), sulcal (i.e., travel) depth, and mean curvature for each cortical brain region. For contextualizing the less common morphometric measures, sulcal depth corresponds to the distance between points on the cortical surface and an outer reference surface that expands across gyri without dipping into sulci, and mean curvature corresponds to the local folding of gyri and sulci (van Essen, 2005; Klein et al., 2017). Global measures were computed by summing or using a weighted average of regional measures to produce total brain volume, total gray matter volume, total white matter volume, total surface area, average cortical thickness, average sulcal depth, and average curvature. Importantly, this processing stream was reproducible, as is akin to other Docker-compliant brain imaging tools (e.g., fMRIPrep; Esteban et al., 2019).

### Statistical analyses

Previous reports by our group have reported on associations between measures of SES and chronic inflammation in a partially overlapping cohort (Jensen et al., 2019a). We also examined this relationship in our cohort by testing for Pearson correlations between chronic inflammation (i.e., frequency of CRP elevations) and maternal education and (log of) income-to-needs.

We addressed our first line of inquiry, whether chronic inflammation relates to brain morphometry, by submitting total and regional volumetric and surface-based measures to Pearson semi-partial correlation analyses controlling brain estimates for age at time of scan and biological sex. Head circumference was also examined for an association with frequency of CRP elevations, but it was not related (p > 0.1). To control for multiple testing for confirmatory analyses, false-discovery rate (FDR) was implemented for volumetric measures of *a priori* selected regions: total brain volume, cerebral gray matter volume, cerebral white matter volume, nucleus accumbens, caudate, putamen, pallidum, thalamus, amygdala, and hippocampus (both hemispheres for basal ganglia and medial temporal lobe regions; Benjamini and Hochberg, 1995). For basal ganglia and medial temporal lobe regions, left and right hemispheres were examined separately to account for the possibility of hemispheric-specific findings, as reported in prior brain imaging studies of inflammation (Graham et al., 2016; Orhun et al., 2018). Consistent with recommendations to refrain from applying confirmatory statistics to results of exploratory analyses (Flournoy et al., 2020) and with previous studies conducting exploratory analyses (e.g., Croarkin et al., 2018), FDR correction was not implemented for exploratory analyses examining surface-based measures (62 parcels for each of the four measures). All associations between chronic inflammation and regional brain morphometry were also re-computed with HAZ and total brain volume added as covariates of no interest. All correlation analyses were computed in Matlab R2016a.

To address the second line of inquiry, whether chronic inflammation mediated relationships between measures of SES and brain morphometry, indirect effects were examined whenever chronic inflammation exhibited a significant (after FDR correction for multiple comparisons; Benjamini and Hochberg, 1995) association with measures of SES and brain morphometry. Mediation models included age at time of scan, biological sex, and HAZ, which has been previously associated with brain morphometry (Turesky et al., 2021). Indirect effects were significant when the 95% confidence intervals (based on 10,000 bootstrapped samples) for its proportion of the total effect did not include 0. Mediation models were implemented using the Mediation package in R. Brain maps were generated using the ggseg() function in R. All code used for statistical analyses and visualizations is housed in an openly available repository at https://github.com/TeddyTuresky/BrainMorphometry_Inflammation_BEANstudy_2021.

### Sensitivity analyses

To determine whether observed brain-inflammation associations were driven by heightened inflammation at a given timepoint, we re-examined relationships between brain structural measures identified in the main analysis using absolute CRP concentrations measured at individual timepoints. As data were non-normally distributed (please see Supplementary Fig. 1), we first applied a log-transform (log([CRP] + 1)), consistent as is common with inflammatory marker data (Arfanakis et al., 2013; Chaban et al., 2020; Ford et al., 2003; Jiang et al., 2014; Wersching et al., 2010; Xie et al., 2019b). We then computed semipartial Spearman correlations controlling for the age at time of scan and biological sex. Continuous measures of CRP concentrations, computed by averaging the log-transformed CRP concentrations, were also submitted to partial correlations to determine whether observed associations were specific to the method for defining inflammation chronicity. Our final sensitivity analyses utilized an independent standard for elevated CRP concentrations. Because pediatric inflammation literature offers widely variable benchmarks for determining elevated CRP concentrations (Järvisalo et al., 2002; Tauman et al., 2004; Taylor et al., 2020), we opted to use the sole benchmark provided by the assay’s manufacturer (i.e., 1 mg/L; Immundiagnostik AG), which reflects medium cardiovascular risk (Pearson et al., 2003). Semipartial brain-behavior correlations for basal ganglia structures were re-computed using this independent benchmark.

## RESULTS

### Chronic inflammation is associated with SES

We first examined the relation between chronic inflammation and measures of socioeconomic status (SES). Frequency of elevations of C-reactive protein (CRP) negatively correlated with maternal education (r = −0.27, p = 0.014, p_FDR_ < 0.05), but not income-to-needs (r = −0.19, p = 0.097).

### Chronic inflammation relates to brain morphometry

We next examined the association between chronic inflammation and estimates of brain volume in several regions selected *a priori*. Frequency of CRP elevations negatively correlated with total brain volume and cerebral white matter volume, but these effects did not remain significant after correction for multiple comparisons. Among the subcortical structures examined, however, frequency of CRP elevations negatively correlated with bilateral caudate, putamen, and pallidum and these effects remained after FDR-correction for multiple comparisons. Brain-inflammation associations for putamen volumes, where effects were strongest, are shown in Figure 1. Except for left pallidum, these effects also remained significant (p < 0.05) when volumes were adjusted for height-for-age (HAZ), an anthropometric marker that has been linked with biological and psychosocial adversity and also to brain structure (Turesky et al., 2021), and total brain volume (Table 1).

**Figure 1.**
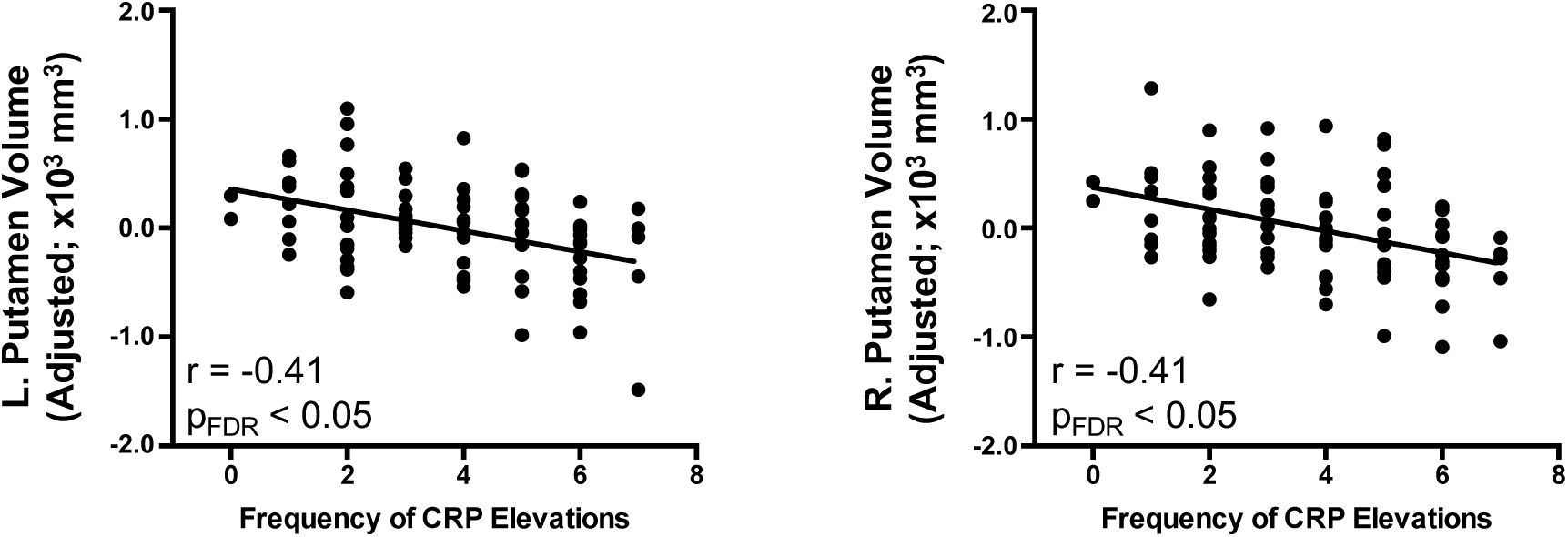
Chronic inflammation predicts bilateral putamen volumes. Chronic inflammation was measured as the frequency of C-reactive protein (CRP) elevations from infancy to age five. Relationships were computed using semipartial correlations with brain measures adjusted for age and biological sex.

**Table 1.**
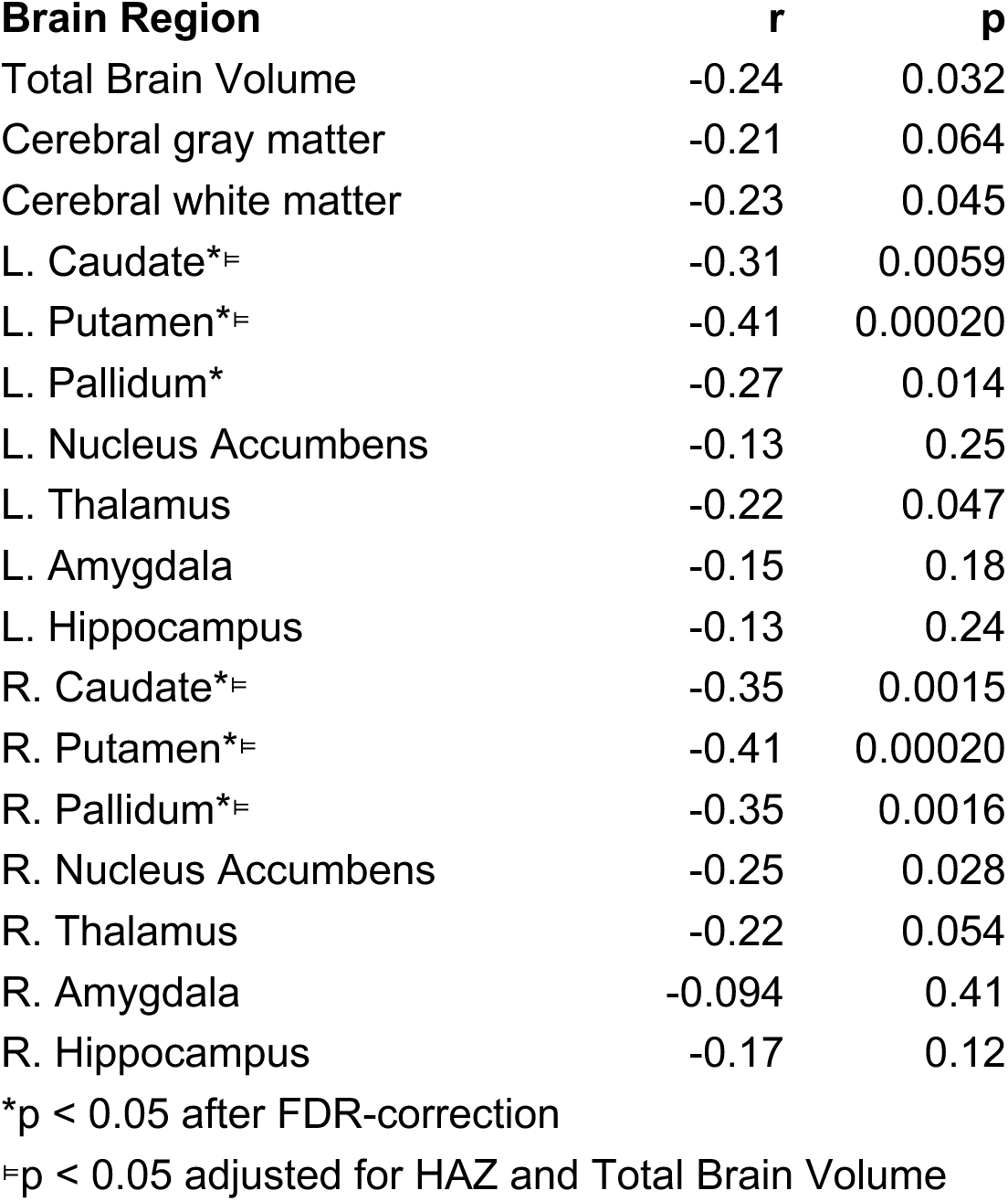
Associations between brain volume and chronic inflammation.

As an exploratory portion of this study, we next examined whether global and regional surface-based measures also related to chronic inflammation. Average sulcal depth was the only global surface-based measure to relate to frequency of CRP elevations (r = −0.24, p = 0.035), but this was prior to correction for multiple comparisons (please see Methods section). Regionally, associations between frequency of CRP elevations and surface-based measures were predominantly for surface area, with fewer associations for cortical thickness, sulcal depth, and mean curvature (Table 2). Surface-based measures in several regions remained significant after controlling for HAZ and total intracranial volume (p < 0.05).

**Table 2.** Associations between brain surface-based measures and chronic inflammation.

| **Brain Region** | **r** | **p** |
| --- | --- | --- |
| *Surface Area* |  |  |
| L. Entorhinal Cortex | -0.29 | 0.0095 |
| L. Parahippocampal Cortex | -0.28 | 0.013 |
| L. Postcentral Gyrus | -0.23 | 0.038 |
| L. Posterior Cingulate Cortex<sup>‡</sup> | -0.33 | 0.0027 |
| L. Precuneus | -0.23 | 0.037 |
| L. Superior Parietal Cortex | -0.26 | 0.020 |
| L. Supramarginal Gyrus | -0.26 | 0.023 |
| R. Fusiform Gyrus | -0.23 | 0.038 |
| R. Parahippocampal Cortex<sup>‡</sup> | -0.28 | 0.013 |
| R. Pars Orbitalis | -0.24 | 0.034 |
| R. Pars Triangularis<sup>‡</sup> | -0.29 | 0.0089 |
| R. Superior Frontal Gyrus | -0.22 | 0.048 |
| R. Insula | -0.27 | 0.016 |
| *Cortical Thickness* |  |  |
| R. Lateral Orbitofrontal Cortex<sup>‡</sup> | -0.23 | 0.045 |
| *Sulcal Depth* |  |  |
| L. Superior Frontal Cortex | -0.25 | 0.025 |
| R. Cuneus<sup>‡</sup> | -0.35 | 0.0014 |
| R. Posterior Cingulate Cortex | -0.26 | 0.020 |
| R. Rostral Middle Frontal Gyrus | -0.28 | 0.011 |
| *Mean Curvature* |  |  |
| L. Paracentral Cortex<sup>‡</sup> | -0.22 | 0.050 |
| L. Posterior Cingulate Cortex<sup>‡</sup> | 0.31 | 0.0048 |
| R. Pericalcarine Cortex | -0.25 | 0.028 |
| <sup>‡</sup>$p < 0.05$ adjusted for HAZ and Total Brain Volume (in addition to age and biological sex already in the model) | | |

### Chronic inflammation mediates the link between maternal education and putamen volume

Previous reports have demonstrated associations between SES and brain morphometry (Noble et al., 2012; Hanson et al., 2013; Luby et al., 2013; Hair et al., 2015; Mackey et al., 2015; Noble et al., 2015; McDermott et al., 2019; Turesky et al., 2021). Our next goal was to determine whether chronic inflammation mediated these or similar links. As a precondition for mediation, the mediator (i.e., frequency of CRP elevations) must be associated with both the predictor (i.e., SES) and outcome (i.e., brain morphometry). Consequently, we examined indirect effects only where frequency of CRP elevations correlated with a measure of SES and a measure of brain morphometry (after FDR correction). Indirect effects are reported significant where 95% confidence intervals (CI) for proportion mediated, based on 10,000 bootstrapped samples, do not include 0. Frequency of CRP elevations mediated the link between maternal education and left (proportion mediated = 0.25, CI [0.0079 0.72], p = 0.044) and right (proportion mediated = 0.21, CI [0.0090 0.57], p = 0.042) putamen (Fig. 2). No other region examined exhibited indirect effects that were significant proportions of their total effect, though right caudate did also exhibit a significant indirect effect (unstandardized effect size = 7.0794, CI [0.055 17.0], p = 0.048).

**Figure 2.**
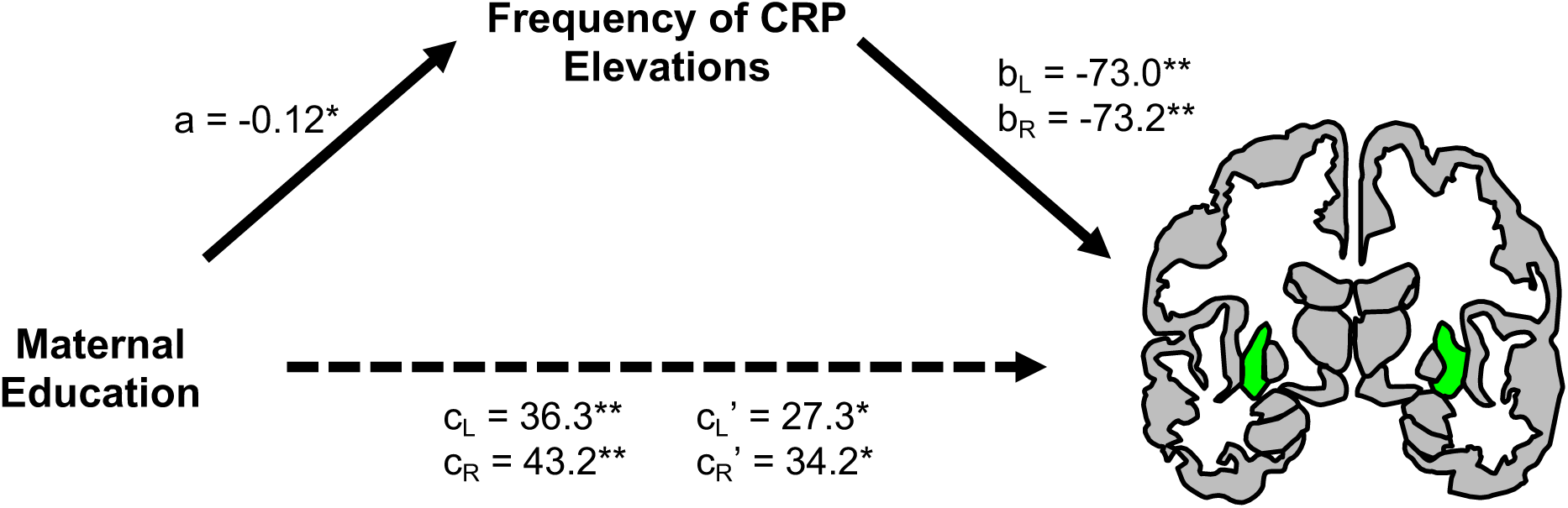
Chronic inflammation mediates an association between maternal education and bilateral putamen volumes (green). Models for indirect effects included age, sex, and HAZ covariates. Unstandardized path coefficients are provided for left (L) and right (R) volumes. *p < 0.05, **p < 0.01

### Sensitivity analysis

In our first sensitivity analysis, regional volumes identified as significant (after FDR correction) in the main analysis were submitted to additional analyses to determine whether effects were driven by heightened CRP levels at a specific time point. Tests of semipartial correlations between absolute concentrations of CRP at individual timepoints and bilateral caudate, putamen, and pallidum volumes showed substantially lower effect sizes overall, with only bilateral putamen and pallidum exhibiting significant (uncorrected) effects and only for CRP measurements made in infancy (Supplementary Table 2).

In our second sensitivity analysis, we examined whether observed effects were specific to the manner in which inflammation chronicity was measured with additional tests of correlation conducted between average (across developmental timepoints) concentrations of CRP and basal ganglia structures identified in the main analysis. Overall, effect sizes were substantially lower using absolute measures of CRP (Supplementary Table 3).

In our third sensitivity analysis, we re-computed semipartial brain-behavior correlations in using an independent benchmark for elevated CRP (1 mg/L). Effect sizes were highly similar compared with the main analysis, with slightly higher values for right putamen and left pallidum and lower values for the other basal ganglia regions (Supplementary Table 4).

## DISCUSSION

Previous work has identified links between inflammation and brain structure and function (Yoon et al., 1997; Duggan et al., 2001; Bø et al., 2003; Hansen-Pupp et al., 2005; Tiberio et al., 2005; Vercellino et al., 2005; Prinster et al., 2006; Capuron et al., 2007; Jefferson et al., 2007; Marsland et al., 2008; Fisher et al., 2008; Capuron et al., 2012; Gianaros et al., 2013; Graham et al., 2018; Rudolph et al., 2018; Sugimoto et al., 2018; Goldstein et al., 2021), but most studies have examined inflammation experienced during a narrow window of time, rather than chronic inflammation (c.f., Xie et al., 2019a). This is an important gap to address as chronic inflammation is prevalent in low-resource areas (Nelson, 2017). We observed that chronic inflammation, specifically, C-reactive protein (CRP) collected at eight time points between birth and five years of age, negatively correlated with gray matter volume in bilateral basal ganglia structures. Further, chronic inflammation partially mediated the relationship between socioeconomic status (SES) and bilateral putamen volumes, suggesting one possible pathway by which SES relates to brain morphometry.

The observed link between chronic inflammation in childhood and volume in basal ganglia structures has a strong foundation in earlier reports on brain-inflammation interactions in adults with inflammatory conditions (Prinster et al., 2006; Westhoff et al., 2019) and adults treated for hepatitis C virus and malignant melanomas with inflammatory cytokines (Capuron et al., 2007, 2012). The specificity to these structures is compelling in light of previous reports showing reduced activity and altered dopamine function in the caudate and putamen with administration of inflammatory cytokine (Capuron et al., 2007, 2012). Broadly, proposed mechanisms point to a disruption in the synthesis and synaptic release of dopamine (Capuron et al., 2012), which reduces synaptic activity, leading to neuronal death (Fricker et al., 2018), and with it, loss of gray matter. Indeed, if dopaminergic systems are susceptible to inflammation, then the caudate and putamen could be particularly affected as they comprise medium spiny neurons with dopamine receptors that receive projections along the nigrostriatal pathway (Alexander and Crutcher, 1990), one of the brain’s chief dopaminergic pathways. Another explanation is that inflammation activates microglia (Sankowski et al., 2015), which in turn can affect neuronal death (Bessis et al., 2007). In support of this, greater numbers of activated microglial cells have been found in putamen, in addition to hippocampus and cerebellum, in individuals with sepsis compared with controls (Westhoff et al., 2019). As the basal ganglia is thought to subserve simple and complex movements (Lehéricy et al., 2006), it is possible that alterations in caudate and putamen structure could also explain the link between chronic inflammation and lower motor performance in early childhood (Jiang et al., 2017; Jensen et al., 2019a). Ultimately, though, future studies are needed to clarify the specific mechanism(s) linking chronic inflammation to brain volume in the basal ganglia.

Associations between chronic inflammation and thalamus, medial temporal lobe structures, and cerebral white matter volume were also hypothesized, based largely on previous studies in infants exposed to elevated inflammatory marker concentrations prenatally (Yoon et al., 1997; Duggan et al., 2001; Hansen-Pupp et al., 2005; Graham et al., 2018) and in clinical populations with characteristically heightened inflammation, such as multiple sclerosis (Compston and Coles, 2008) and sepsis (Orhun et al., 2020). However, chronic inflammation did not exhibit significant relations to thalamus or medial temporal lobe volumes, and the relation to cerebral white matter volume exhibited a markedly lower effect size, which was only significant *prior* to correction for multiple comparisons. Interestingly, though, the relatively low effect size is fairly consistent with reports examining relations between inflammation and mean fractional anisotropy (Gianaros et al., 2013) and neurodevelopmental outcomes (Jiang et al., 2014, 2017), suggesting that inflammation may not explain a substantial portion of the variance in global white matter organization or behavioral measures. It is also worth noting that neither CRP (the inflammatory marker assayed for the present study) nor any other inflammatory marker related to total white matter volume (after controlling for total intracranial volume) in a cross-sectional study in adults (Jefferson et al., 2007). It is also possible that results observed in the current study diverged from the literature in these regions because different inflammatory markers do not necessarily relate to the brain in the same manner; e.g., interleukin 6 and tumor necrosis factor alpha related to total brain volume, but monocyte chemoattractant protein-1 did not (Jefferson et al., 2007). Similarly, most studies examining brain-inflammation relationships did so by assaying inflammatory cytokines (e.g., interleukins), whereas CRP is synthesized downstream in an inflammatory response. Consequently, it may behoove future developmental cognitive neuroscience studies examining chronic inflammation to also measure cytokine concentrations across developmental stages.

We also observed associations between chronic inflammation and surface-based brain measures. While few spatial consistencies were observed across measures—e.g., left posterior cingulate cortex was the only brain area related to chronic inflammation on multiple measures (surface area and sulcal depth)—all correlations that were observed were in the same direction (i.e., negative). Additionally, global mean sulcal depth was associated with chronic inflammation, which is consistent with literature showing this measure’s sensitivity to other early adversities, such as preterm brain injury (Shimony et al., 2016; albeit prior to controlling for total intracranial volume) and exposure to prenatal alcohol (De Guio et al., 2014). Nevertheless, these results should be viewed with caution; due to the exploratory approach we implemented for surface-based measures, we did not tether results here to confirmatory statistics (e.g., with corrections for multiple comparisons; Flournoy et al., 2020). Consequently, it may behoove future studies with larger, independent samples to use the surface-based findings presented here as a starting-point for more hypothesis-driven approaches and more rigorous statistical thresholding.

In a broader context, the observation that chronic inflammation mediates a significant portion of the relationship between SES and bilateral putamen volume addresses a consistent gap in the multitudes of reports linking SES to brain structure and function without intermediate, proximal factors to explain this association (Farah, 2017). As such, it suggests that chronic inflammation may join stress (Luby et al., 2013; Merz et al., 2019), caregiving quality (Luby et al., 2013), home linguistic environment (Merz et al., 2020), and diminished growth (which serves as a proxy for malnutrition and other biological and psychosocial hazards; Turesky et al., 2021), as another proximal factor for ‘embedding’ low SES in brain development (Jensen et al., 2017). Future studies will be needed to identify other candidate risk factors associated with SES (e.g., toxins, cognitive stimulation; Farah, 2017).

A final contribution of this study is that it does not rely on data from white participants from high-resource countries, which differentiates it from most other scientific studies (Cell Editorial Team, 2020). In contrast, recruited children from the low-resource country Bangladesh, and in so doing begins to address the racial and socioeconomic inequity endemic to developmental cognitive neuroscience.

Nevertheless, this study had four main limitations. The first three stem from the way chronic inflammation was measured, namely, as the frequency of peripheral CRP concentration elevations. First, while this method has been previously employed (Naylor et al., 2015; Jiang et al., 2017; Jensen et al., 2019b; Xie et al., 2019b; Bach et al., 2022), it is an estimate calculated from only eight measurements of CRP concentration across five years and using a cohort-specific threshold for separating elevated from normative CRP levels, rather than an independent standard. While our first sensitivity analysis, showing attenuated effects when inflammation was measured at individual timepoints, suggests that the observed findings in the main analysis were not driven by heightened inflammation at any given timepoint(s), our second and third sensitivity analyses using average (across developmental time points) CRP concentrations and an independent CRP concentration benchmark suggest specificity to the particular method of measuring inflammatory chronicity, but not necessarily the specific benchmark value. Second, CRP is one of numerous inflammatory markers, which were not examined as part of the calculation of chronic inflammation in this study. Although other inflammatory markers were assayed in these children, CRP was measured more frequently than any other marker. As with prior studies (Jiang et al., 2017; Jensen et al., 2019b; Xie et al., 2019b; Bach et al., 2022), we opted for frequency of repeated sampling over a comprehensive estimate of inflammation with fewer repeated measures. Third, while inflammatory markers can penetrate the central nervous system (Sankowski et al., 2015), CRP was assayed from the periphery. Fourth, the analyses presented here account for age, biological sex, and anthropometry. However, the low-resource environment in which the children in this study grow up exposes them to myriad severe biological and psychosocial risk factors (Jensen et al., 2017; Nelson, 2017), for which the present study does not control. Future studies with larger sample sizes and gathering more inflammatory marker data, particularly that which exceeds the independent benchmark, will be needed to address these limitations.

## CONCLUSION

This study examined chronic inflammation and brain morphometry in children reared in an extremely impoverished area of Bangladesh, a population that is highly underrepresented in developmental cognitive research. Our findings indicate that chronic inflammation relates to brain structure and partially mediates an association between socioeconomic status and brain volume in bilateral putamen in children growing up in extreme poverty in Bangladesh. These findings suggest that chronic inflammation may represent one proximal factor through which poverty derails typical brain development. Future intervention studies that reduce chronic inflammation in children growing up in low-resource settings will be needed to test this potential causality, but this study represents an important first step in considering how chronic inflammation might impact brain development.

## ACKNOWLEDGEMENTS

Funding for this work was provided by research grants from the Bill & Melinda Gates Foundation to CAN [OPP1111625] and WAP [OPP1017093], from the Henske Foundation to WAP, from NIAID (R01 AI043596-17) to WAP, and from the Harvard Brain Initiative Transitions Program to TKT. We are grateful to the families who participated in the study, the staff at The International Centre for Diarrhoeal Disease Research (ICDDR,B) who undertook and completed data collection, and Uma Nayak and Rachel Kwon for organizing the non-MRI measures. Finally, we thank the Harvard Catalyst Biostatistical Consulting program for guidance on reporting mediations.

## Supplementary Tables

**Supplementary Table 1.** Descriptive statistics for concentrations of C-reactive protein.

| Timepoint<br>(weeks) | Median<br>(mg/L) | Minimum<br>(mg/L) | Maximum<br>(mg/L) |
| --- | --- | --- | --- |
| 6 | 0.085 | 0 | 7.37 |
| 18 | 0.5 | 0 | 33.42 |
| 40 | 1.725 | 0 | 72.58 |
| 53 | 0.82 | 0 | 28.88 |
| 104 | 1.2 | 0 | 62.87 |
| 156 | 0.55 | 0.4 | 110.05 |
| 207 | 0.58 | 0.04 | 58.34 |
| 260 | 1.365 | 0 | 57.59 |

**Supplementary Table 2.**
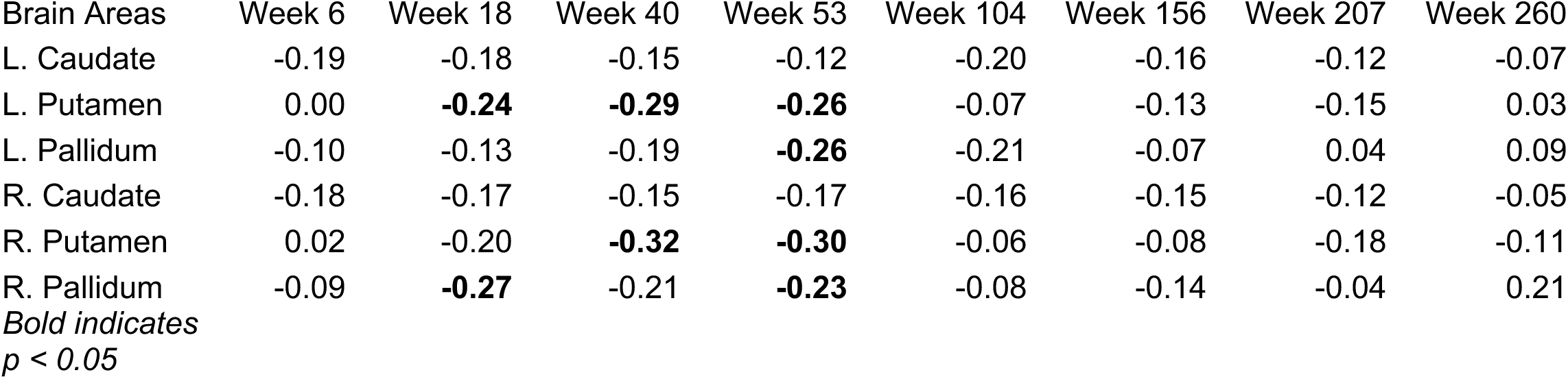
Associations between basal ganglia volumes and inflammation across developmental time points.

**Supplementary Table 3.** Associations between basal ganglia volumes and average inflammation across developmental time points.

| Brain Areas | r | p |
| --- | --- | --- |
| L. Caudate | -0.23 | 4.4E-02 |
| L. Putamen | -0.22 | 5.7E-02 |
| L. Pallidum | -0.19 | 9.7E-02 |
| R. Caudate | -0.24 | 3.2E-02 |
| R. Putamen | -0.28 | 1.2E-02 |
| R. Pallidum | -0.19 | 1.0E-01 |

**Supplementary Table 4.**
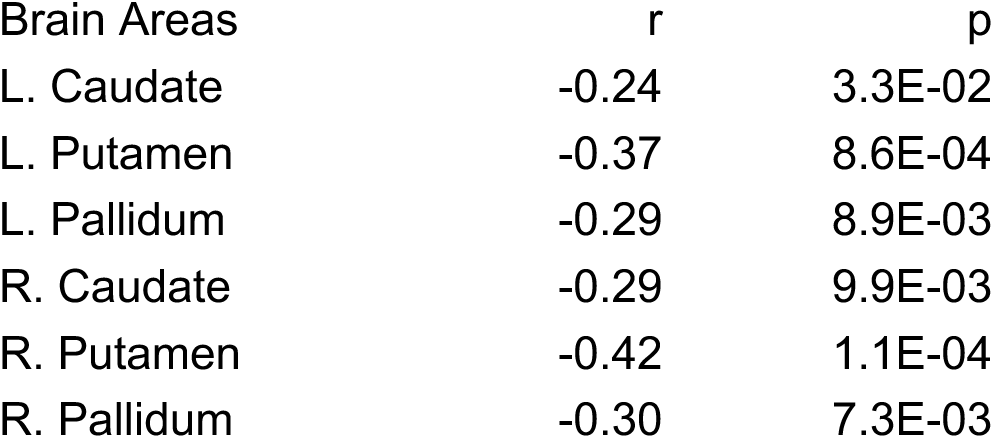
Associations between basal ganglia volumes and chronic inflammation with elevations measured as [CRP] > 1 mg/L.

## Supplementary Figures

**Supplementary Figure 1.**
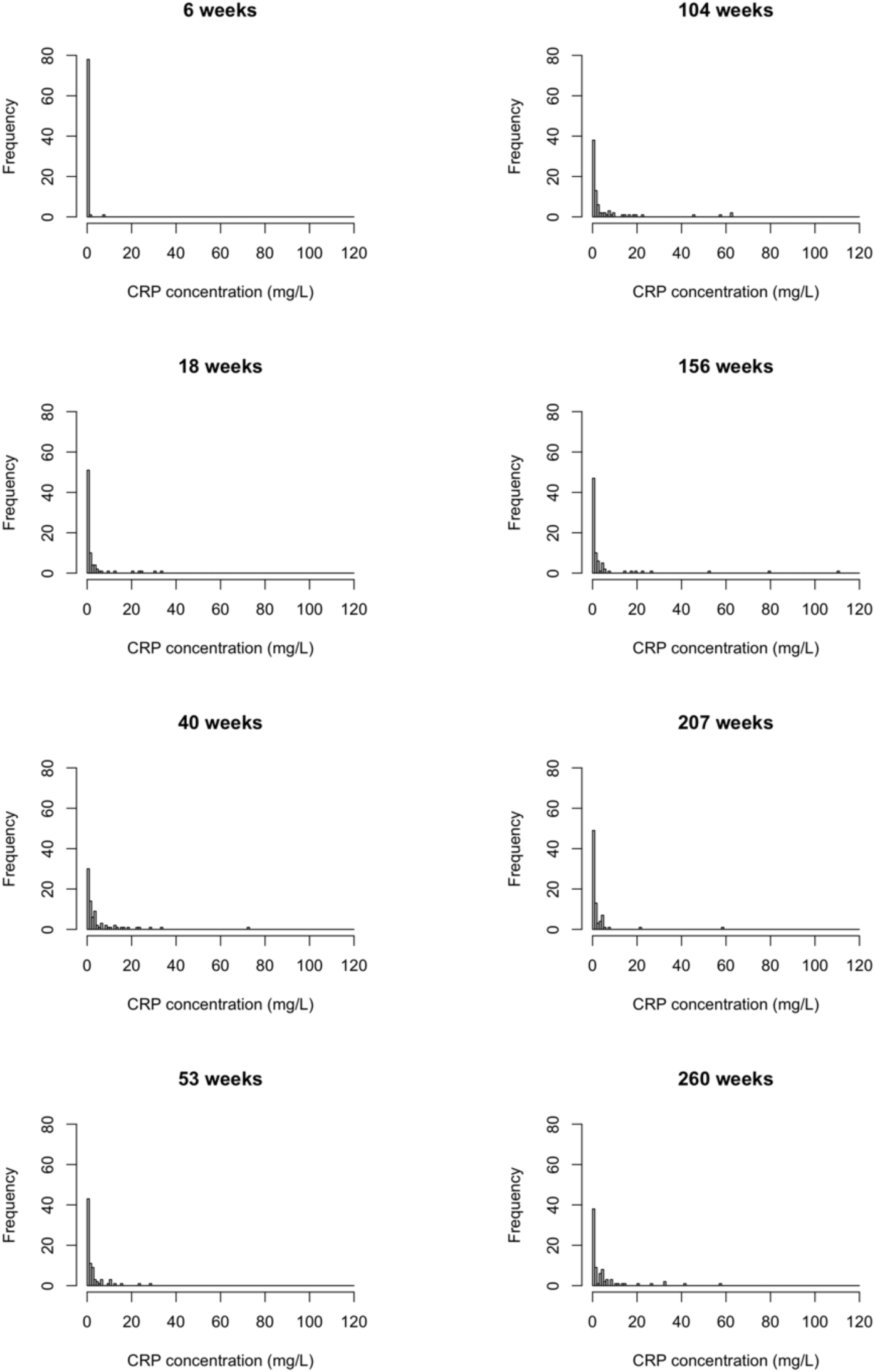
Distribution of C-reactive protein (CRP) concentrations. Histograms depict the distribution of CRP concentrations across eight developmental time points.

## REFERENCES

Adler, N., Boyce, T., Chesney, M., Cohen, S., Folkman, S., Kahn, R., Syme, S., 1994. Socioeconomic status and health: the challenge of the gradient. American Psychologist 49, 15–24. 10.1021/jo901279g

Alexander, G.E., Crutcher, M.D., 1990. Functional architecture of basal ganglia circuits: neural substrates of parallel processing. Trends Neurosci 13, 266–71.

Arfanakis, K., Fleischman, D.A., Grisot, G., Barth, C.M., Varentsova, A., Morris, M.C., Barnes, L.L., Bennett, D.A., 2013. Systemic Inflammation in Non-Demented Elderly Human Subjects: Brain Microstructure and Cognition. PLoS One 8. 10.1371/journal.pone.0073107

Atalabi, O.M., Lagunju, I.A., Tongo, O.O., Akinyinka, O.O., 2010. Cranial magnetic resonance imaging findings in Kwashiorkor. International journal of neuroscience 120, 23–27. 10.3109/00207450903315727

Bach, A.M., Xie, W., Piazzoli, L., Jensen, S.K.G., Afreen, S., Haque, R., Petri, W.A., Nelson, C.A., 2022. Systemic inflammation during the first year of life is associated with brain functional connectivity and future cognitive outcomes. Dev Cogn Neurosci 53. 10.1016/j.dcn.2021.101041

Benjamini, Y., Hochberg, Y., 1995. Controlling the false discovery rate : a practical and powerful approach to multiple testing. J.R. Statistics 57, 289–300.

Bessis, A., Bechade, C., Bernard, D., Roumier, A., 2007. Microglial control of neuronal death and synaptic properties ALAIN. Glia 55, 233–238. 10.1002/glia

Betancourt, L.M., Avants, B., Farah, M.J., Brodsky, N.L., Wu, J., Ashtari, M., Hurt, H., 2016. Effect of socioeconomic status (SES) disparity on neural development in female African-American infants at age 1 month. Dev Sci 19, 947–956. 10.1111/desc.12344

Bhutta, Z.A., Guerrant, R.L., Nelson, C.A., 2017. Neurodevelopment, Nutrition, and Inflammation: The Evolving Global Child Health Landscape. Pediatrics 139, S12–S22. 10.1542/peds.2016-2828D

Bø, L., Vedeler, C.A., Nyland, H.I., Trapp, B.D., Mørk, S.J., 2003. Subpial demyelination in the cerebral cortex of multiple sclerosis patients. J Neuropathol Exp Neurol 62, 723–732. 10.1093/jnen/62.7.723

Bosmann, M., Ward, P.A., 2013. The inflammatory response in sepsis. Trends Immunol 34, 129–136. 10.1016/j.it.2012.09.004

Brito, N.H., Fifer, W.P., Myers, M.M., Elliott, A.J., Noble, K.G., 2016. Associations among family socioeconomic status, EEG power at birth, and cognitive skills during infancy. Dev Cogn Neurosci 19, 144–151. 10.1016/j.dcn.2016.03.004

Brito, N.H., Noble, K.G., 2014. Socioeconomic status and structural brain development. Front Neurosci 8, 1–12. 10.3389/fnins.2014.00276

Calderón-Garcidueñas, L., Engle, R., Antonieta Mora-Tiscareño, A.M., Styner, M., Gómez-Garza, G., Zhu, H., Jewells, V., Torres-Jardón, R., Romero, L., Monroy-Acosta, M.E., Bryant, C., González-González, L.O., Medina-Cortina, H., D’Angiulli, A., 2011. Exposure to severe urban air pollution influences cognitive outcomes, brain volume and systemic inflammation in clinically healthy children. Brain Cogn 77, 345–355. 10.1016/j.bandc.2011.09.006

Capuron, L., Pagnoni, G., Demetrashvili, M.F., Lawson, D.H., Fornwalt, F.B., Woolwine, B., Berns, G.S., Nemeroff, C.B., Miller, A.H., 2007. Basal ganglia hypermetabolism and symptoms of fatigue during interferon-α therapy. Neuropsychopharmacology 32, 2384– 2392. 10.1038/sj.npp.1301362

Capuron, L., Pagnoni, G., Drake, D.F., Woolwine, B.J., Spivey, J.R., Crowe, R.J., Votaw, J.R., Goodman, M.M., Miller, A.H., 2012. Dopaminergic mechanisms of reduced basal ganglia responses to hedonic reward during interferon alfa administration. Arch Gen Psychiatry 69, 1044–1053. 10.1001/archgenpsychiatry.2011.2094

Cell Editorial Team, 2020. Science Has a Racism Problem. Cell 181, 1443–1444. 10.1016/j.cell.2020.06.009

Chaban, V., Clarke, G.J.B., Skandsen, T., Islam, R., Einarsen, C.E., Vik, A., Damås, J.K., Mollnes, T.E., Håberg, A.K., Pischke, S.E., 2020. Systemic Inflammation Persists the First Year after Mild Traumatic Brain Injury: Results from the Prospective Trondheim Mild Traumatic Brain Injury Study. J Neurotrauma 37, 2120–2130. 10.1089/neu.2019.6963

Chiang, J.J., Taylor, S.E., Bower, J.E., 2015. Early adversity, neural development, and inflammation. Dev Psychobiol 57, 887–907. 10.1002/dev.21329

Compston, A., Coles, A., 2008. Multiple sclerosis. The Lancet 372, 1502–1517. 10.1016/S0140-6736(08)61620-7

Croarkin, P.E., Nakonezny, P.A., Deng, Z. de, Romanowicz, M., Voort, J.L.V., Camsari, D.D., Schak, K.M., Port, J.D., Lewis, C.P., 2018. High-frequency repetitive TMS for suicidal ideation in adolescents with depression. J Affect Disord 239, 282–290. 10.1016/j.jad.2018.06.048

De Guio, F., Mangin, J.F., Rivière, D., Perrot, M., Molteno, C.D., Jacobson, S.W., Meintjes, E.M., Jacobson, J.L., 2014. A study of cortical morphology in children with fetal alcohol spectrum disorders. Hum Brain Mapp 35, 2285–2296. 10.1002/hbm.22327

de Onis, M., Garza, C., Onyango, A., Martorell, R., 2006. WHO child growth standards. Acta Paediatr 95, 5–6. 10.1007/s00431-009-1041-x

de Onis, M., Garza, C., Victora, C.G., Bhan, M.K., Norum, K.R., 2004. The WHO Multicentre Growth Reference Study (MGRS): Rationale, planning, and implementation. Food Nutr Bull 25, S3–S89. 10.1177/15648265040251S103

Diamond, B., Honig, G., Mader, S., Brimberg, L., Volpe, B.T., 2013. Brain-reactive antibodies and disease. Annu Rev Immunol 31, 345–385. 10.1146/annurev-immunol-020711-075041

Donowitz, J.R., Cook, H., Alam, M., Tofail, F., Kabir, M., Colgate, E.R., Carmolli, M.P., Kirkpatrick, B.D., Nelson, C.A., Ma, J.Z., Haque, R., Petri, W.A., 2018. Role of maternal health and infant inflammation in nutritional and neurodevelopmental outcomes of two-year-old Bangladeshi children. PLoS Negl Trop Dis 12, 1–20. 10.1371/journal.pntd.0006363

Duggan, P.J., Maalouf, E.F., Watts, T.L., Sullivan, M.H.F., Counsell, S.J., Allsop, J., Al-Nakib, L., Rutherford, M.A., Battin, M., Roberts, I., Edwards, A.D., 2001. Intrauterine T-cell activation and increased proinflammatory cytokine concentrations in preterm infants with cerebral lesions. Lancet 358, 1699–1700. 10.1016/S0140-6736(01)06723-X

Duncan, G.J., Magnuson, K., Votruba-Drzal, E., 2017. Moving beyond Correlations in Assessing the Consequences of Poverty. Annu Rev Psychol 68, 413–434. 10.1146/annurev-psych-010416-044224

El-Sherif, A.M., Babrs, G.M., Ismail, A.M., 2012. Cranial Magnetic Resonance Imaging (MRI) Changes in Severely Malnourished Children before and after Treatment. Life Sci J 9, 738– 742.

Esteban, O., Markiewicz, C.J., Blair, R.W., Moodie, C.A., Isik, A.I., Erramuzpe, A., Kent, J.D., Goncalves, M., DuPre, E., Snyder, M., Oya, H., Ghosh, S.S., Wright, J., Durnez, J., Poldrack, R.A., Gorgolewski, K.J., 2019. fMRIPrep: a robust preprocessing pipeline for functional MRI. Nat Methods 16, 111–116. 10.1038/s41592-018-0235-4

Farah, M.J., 2018. Socioeconomic status and the brain: Prospects for neuroscience-informed policy. Nat Rev Neurosci 19, 428–438. 10.1038/s41583-018-0023-2

Farah, M.J., 2017. The Neuroscience of Socioeconomic Status: Correlates, Causes, and Consequences. Neuron 96, 56–71. 10.1016/j.neuron.2017.08.034

Favrais, G., Looij, Y. van de, Fleiss, B., Ramanantsoa, N., Bonnin, P., Stoltenburg-didinger, G., Lacaud, A., Saliba, E., Dammann, O., Gallego, J., Sizonenko, S., Hagberg, H., Lelievre, V., Gressens, P., 2011. Systemic Inflammation Disrupts the Developmental Program of White Matter. Ann Neurol 70, 550–65. 10.1002/ana.22489

Fischl, B., Salat, D., Busa, E., Albert, M., Dieterich, M., Haselgrove, C., van der Kouwe, A., Killiany, R., Kennedy, D., Klaveness, S., Montillo, A., Makris, N., Rosen, B., Dale, A., 2002. Whole Brain Segmentation: Automated Labeling of Neuroanatomical Structures in the Human Brain. Neuron 33, 341–355.

Fisher, E., Lee, J.C., Nakamura, K., Rudick, R.A., 2008. Gray matter atrophy in multiple sclerosis: A longitudinal study. Ann Neurol 64, 255–265. 10.1002/ana.21436

Fitzpatrick, Z., Frazer, G., Ferro, A., Clare, S., Bouladoux, N., Ferdinand, J., Tuong, Z.K., Negro-Demontel, M.L., Kumar, N., Suchanek, O., Tajsic, T., Harcourt, K., Scott, K., Bashford-Rogers, R., Helmy, A., Reich, D.S., Belkaid, Y., Lawley, T.D., McGavern, D.B., Clatworthy, M.R., 2020. Gut-educated IgA plasma cells defend the meningeal venous sinuses. Nature 587, 472–476. 10.1038/s41586-020-2886-4

Flournoy, J.C., Vijayakumar, N., Cheng, T.W., Cosme, D., Flannery, J.E., Pfeifer, J.H., 2020. Improving practices and inferences in developmental cognitive neuroscience. Dev Cogn Neurosci 45, 100807. 10.1016/j.dcn.2020.100807

Ford, E., Giles, W., Myers, G., Rifai, N., Ridker, P., Mannino, D., 2003. C-reactive Protein Concentration Distribution among US Children and Young Adults: Findings from the National Health and Nutrition Examination Survey, 1999–2000. Clin Chem 49, 1353–1357.

Fricker, M., Tolkovsky, A.M., Borutaite, V., Coleman, M., Brown, G.C., 2018. Neuronal cell death. Physiol Rev 98, 813–880. 10.1152/physrev.00011.2017

Gao, H.M., Jiang, J., Wilson, B., Zhang, W., Hong, J.S., Liu, B., 2002. Microglial activation-mediated delayed and progressive degeneration of rat nigral dopaminergic neurons: Relevance to Parkinson’s disease. J Neurochem 81, 1285–1297. 10.1046/j.1471-4159.2002.00928.x

Gianaros, P.J., Marsland, A.L., Sheu, L.K., Erickson, K.I., Verstynen, T.D., 2013. Inflammatory Pathways Link Socioeconomic Inequalities to White Matter Architecture. Cerebral Cortex 23, 2058–2071. 10.1093/cercor/bhs191

Goldstein, J.M., Cohen, J.E., Mareckova, K., Holsen, L., Whitfield-Gabrieli, S., Gilman, S.E., Buka, S.L., Hornig, M., 2021. Impact of prenatal maternal cytokine exposure on sex differences in brain circuitry regulating stress in offspring 45 years later. Proc Natl Acad Sci U S A 118, 1–8. 10.1073/pnas.2014464118

Graham, A.M., Buss, C., Rasmussen, J.M., Rudolph, M.D., Demeter, D. V., Gilmore, J.H., Styner, M., Entringer, S., Wadhwa, P.D., Fair, D.A., 2016. Implications of newborn amygdala connectivity for fear and cognitive development at 6-months-of-age. Dev Cogn Neurosci 18, 12–25. 10.1016/j.dcn.2015.09.006

Graham, A.M., Rasmussen, J.M., Rudolph, M.D., Heim, C.M., Gilmore, J.H., Styner, M., Potkin, S.G., Entringer, S., Wadhwa, P.D., Fair, D.A., Buss, C., 2018. Maternal Systemic Interleukin-6 During Pregnancy Is Associated With Newborn Amygdala Phenotypes and Subsequent Behavior at 2 Years of Age. Biol Psychiatry 83, 109–119. 10.1016/j.biopsych.2017.05.027

Grantham-McGregor, S., Cheung, Y.B., Cueto, S., Glewwe, P., Richter, L., Strupp, B., International, C.D.S.G., 2007. Child development in developing countries. Lancet 369, 60– 70.

Gunston, G.D., Burkimsher, D., Malan, H., Sive, A.A., 1992. Reversible cerebral shrinkage in kwashiorkor: an MRI study. Arch Dis Child 67, 1030–1032.

Hair, N.L., Hanson, J.L., Wolfe, B.L., Pollak, S.D., 2015. Association of child poverty, brain development, and academic achievement. JAMA Pediatr 169, 822–829. 10.1001/jamapediatrics.2015.1475

Hansen-Pupp, I., Harling, S., Berg, A.C., Cilio, C., Hellström-Westas, L., Ley, D., 2005. Circulating interferon-gamma and white matter brain damage in preterm infants. Pediatr Res 58, 946–952. 10.1203/01.PDR.0000182592.76702.E8

Hanson, J.L., Hair, N., Shen, D.G., Shi, F., Gilmore, J.H., Wolfe, B.L., Pollak, S.D., 2013. Family poverty affects the rate of human infant brain growth. PLoS One 8. 10.1371/journal.pone.0080954

Hazin, A.N., Alves, J.G.B., Falbo, A.R., 2007. The myelination process in severely malnourished children: MRI findings. International Journal of Neuroscience 117, 1209–1214. 10.1080/00207450600934945

Hoff, E., Laursen, B., Tardif, T., 2002. Socioeconomic Status and Parenting, in: Bornstein, M. (Ed.), Handbook of Parenting: Biology and Ecology of Parenting. Lawrence Erlbaum Associates Publishers, pp. 231–252.

Järvisalo, M.J., Harmoinen, A., Hakanen, M., Paakkunainen, U., Viikari, J., Hartiala, J., Lehtimäki, T., Simell, O., Raitakari, O.T., 2002. Elevated serum C-reactive protein levels and early arterial changes in healthy children. Arterioscler Thromb Vasc Biol 22, 1323– 1328. 10.1161/01.ATV.0000024222.06463.21

Jefferson, A., Massaro, J., Wolf, P., Seshadri, S., Au, R., Vasan, R., Larson, M., Meigs, J., Keaney, J., Lipinska, I., Kathiresan, S., Benjamin, E., DeCarli, C., 2007. Inflammatory biomarkers are associated with total brain volume The Framingham Heart Study. Neurology 68, 1032–1039.

Jensen, S., Berens, A., Nelson, C., 2017. Effects of poverty on interacting biological systems underlying child development. Lancet Child Adolesc Health 1, 225–239. 10.1016/S2352-4642(17)30024-X

Jensen, S., Kumar, S., Xie, W., Tofail, F., Haque, R., Petri, W., Nelson, C., 2019a. Neural correlates of early adversity among Bangladeshi infants. Sci Rep 9, 1–10. 10.1038/s41598-019-39242-x

Jensen, S., Tofail, F., Haque, R., Petri, W., Nelson, C., 2019b. Child development in the context of biological and psychosocial hazards among poor families in Bangladesh. PLoS One 14, 1–17. 10.17605/OSF.IO/X2MC7.Funding

Jerry Silver, Martin E. Schwab, and P.G.P., 2015. Central Nervous System Regenerative Failure : Cold Spring Harb Perspect Biol 7, a020602.

Jiang, N., Cowan, M., Moonah, S., Petri, W., 2018. The impact of systemic inflammation on neurodevelopment. Trends Mol Med 24, 794–804.

Jiang, N.M., Tofail, F., Ma, J.Z., Haque, R., Kirkpatrick, B., Nelson, C.A., Petri, W.A., 2017. Early life inflammation and neurodevelopmental outcome in Bangladeshi infants growing up in adversity. American Journal of Tropical Medicine and Hygiene 97, 974–979. 10.4269/ajtmh.17-0083

Jiang, N.M., Tofail, F., Moonah, S.N., Scharf, R.J., Taniuchi, M., Ma, J.Z., Hamadani, J.D., Gurley, E.S., Houpt, E.R., Azziz-Baumgartner, E., Haque, R., Petri, W.A., 2014. Febrile illness and pro-inflammatory cytokines are associated with lower neurodevelopmental scores in Bangladeshi infants living in poverty. BMC Pediatr 14, 1–9. 10.1186/1471-2431-14-50

John, C.C., Black, M.M., Nelson, C.A., 2017. Neurodevelopment: The Impact of Nutrition and Inflammation During Early to Middle Childhood in Low-Resource Settings. Pediatrics 139, S59–S71. 10.1542/peds.2016-2828H

Kamata, M., Higuchi, H., Yoshimoto, M., Yoshida, K., Shimizu, T., 2000. Effect of single intracerebroventricular injection of a-interferon on monoamine concentrations in the rat brain, European Neuropsychopharmacology.

Klein, A., Ghosh, S.S., Bao, F.S., Giard, J., Häme, Y., Stavsky, E., Lee, N., Rossa, B., Reuter, M., Chaibub Neto, E., Keshavan, A., 2017. Mindboggling morphometry of human brains, PLoS Computational Biology. 10.1371/journal.pcbi.1005350

Klein, A., Tourville, J., 2012. 101 labeled brain images and a consistent human cortical labeling protocol. Front Neurosci. 10.3389/fnins.2012.00171

Kumar, N., Goyal, S., Tiwari, K., Goyal, V., Meena, P., 2016. Neuroimaging (MRI) in Children with Microcephaly and Severe Acute Malnutrition. International Journal of Medical Pediatrics and Oncology 2, 15–19.

Kutlesic, V., Brewinski Isaacs, M., Freund, L.S., Hazra, R., Raiten, D.J., 2017. Executive Summary: Research Gaps at the Intersection of Pediatric Neurodevelopment, Nutrition, and Inflammation in Low-Resource Settings. Pediatrics 139, S1–S11. 10.1542/peds.2016-2828C

Lawson, G.M., Duda, J.T., Avants, B.B., Wu, J., Farah, M.J., 2013. Associations between children’s socioeconomic status and prefrontal cortical thickness. Dev Sci 16, 641–652. 10.1111/desc.12096.Associations

Lebel, C., Walton, M., Letourneau, N., Giesbrecht, G.F., Kaplan, B.J., Dewey, D., 2016. Prepartum and Postpartum Maternal Depressive Symptoms Are Related to Children’s Brain Structure in Preschool. Biol Psychiatry 80, 859–868. 10.1016/j.biopsych.2015.12.004

Lehéricy, S., Bardinet, E., Tremblay, L., van de Moortele, P.-F., Pochon, J.-B., Dormont, D., Kim, D.-S., Yelnik, J., Ugurbil, K., 2006. Motor control in basal ganglia circuits using fMRI and brain atlas approaches. Cerebral cortex 16, 149–61. 10.1093/cercor/bhi089

Lelijveld, N., Jalloh, A.A., Kampondeni, S.D., Seal, A., Wells, J.C., Goyheneix, M., Chimwezi, E., Mallewa, M., Nyirenda, M.J., Heyderman, R.S., Kerac, M., 2019. Brain MRI and cognitive function seven years after surviving an episode of severe acute malnutrition in a cohort of Malawian children. Public Health Nutr 22, 1406–1414. 10.1017/S1368980018003282

Luby, J., Belden, A., Botteron, K., Marrus, N., Harms, M.P., Babb, C., Nishino, T., Barch, D., 2013. The effects of poverty on childhood brain development: The mediating effect of caregiving and stressful life events. JAMA Pediatr 167, 1135–1142. 10.1001/jamapediatrics.2013.3139

Mackey, A., Finn, A., Leonard, J., Jacoby-Senghor, D., West, M., Gabrieli, C., Gabrieli, J., 2015. Neuroanatomical correlates of the income-achievement gap. Psychol Sci 26, 925–933. 10.1177/0956797615572233.Neuroanatomical

Marsland, A.L., Gianaros, P.J., Abramowitch, S.M., Manuck, S.B., Hariri, A.R., 2008. Interleukin-6 Covaries Inversely with Hippocampal Grey Matter Volume in Middle-Aged Adults. Biol Psychiatry 64, 484–490. 10.1016/j.biopsych.2008.04.016

McDermott, C.L., Seidlitz, J., Nadig, X.A., Liu, S., Clasen, L.S., Blumenthal, J.D., Reardon, P.K., Franc, X., Greenstein, D., Patel, X.R., Chakravarty, M.M., Lerch, J.P., Raznahan, X.A., 2019. Longitudinally mapping childhood socioeconomic status associations with cortical and subcortical morphology. Journal of Neuroscience 39, 1365–1373.

McLaughlin, K.A., Sheridan, M.A., Winter, W., Fox, N.A., Zeanah, C.H., Nelson, C.A., 2014. Widespread reductions in cortical thickness following severe early-life deprivation: A neurodevelopmental pathway to attention-deficit/hyperactivity disorder. Biol Psychiatry 76, 629–638. 10.1016/j.biopsych.2013.08.016

Merz, E., Maskus, E., Melvin, S., He, X., Noble, K., 2020. Socioeconomic disparities in language input are associated with children’s language-related brain structure and reading skills. Child Dev 91, 846–860.

Merz, E.C., Desai, P.M., Maskus, E.A., Melvin, S.A., Rehman, R., Torres, S.D., Meyer, J., He, X., Noble, K.G., 2019. Socioeconomic Disparities in Chronic Physiologic Stress Are Associated With Brain Structure in Children. Biol Psychiatry 86, 921–929. 10.1016/j.biopsych.2019.05.024

Merz, E.C., Tottenham, N., Noble, K.G., 2018. Socioeconomic Status, Amygdala Volume, and Internalizing Symptoms in Children and Adolescents. Journal of Clinical Child and Adolescent Psychology 47, 312–323. 10.1080/15374416.2017.1326122

Moreau, G., Ramakrishnan, G., Cook, H., Fox, T., Nayak, U., Kumar, S., Ma, J., Colgate, E., Kirkpatrick, B., Nelson, C., Haque, R., Petri, W., 2019. Childhood Growth and Neurocognition are Associated with Distinct Sets of Metabolites Article. EBioMedicine in press.

Naylor, C., Lu, M., Haque, R., Mondal, D., Buonomo, E., Nayak, U., Mychaleckyj, J.C., Kirkpatrick, B., Colgate, R., Carmolli, M., Dickson, D., van der Klis, F., Weldon, W., Steven Oberste, M., Ma, J.Z., Petri, W.A., The PROVIDE study teams, 2015. Environmental Enteropathy, Oral Vaccine Failure and Growth Faltering in Infants in Bangladesh. EBioMedicine 2, 1759–1766. 10.1016/j.ebiom.2015.09.036

Nelson, C.A., 2017. Hazards to Early Development: The Biological Embedding of Early Life Adversity. Neuron 96, 262–266. 10.1016/j.neuron.2017.09.027

Noble, K.G., Houston, S.M., Brito, N.H., Bartsch, H., Kan, E., Kuperman, J.M., Akshoomoff, N., Amaral, D.G., Bloss, C.S., Libiger, O., Schork, N.J., Murray, S.S., Casey, B.J., Chang, L., Ernst, T.M., Frazier, J.A., Gruen, J.R., Kennedy, D.N., Van Zijl, P., Mostofsky, S., Kaufmann, W.E., Kenet, T., Dale, A.M., Jernigan, T.L., Sowell, E.R., 2015. Family income, parental education and brain structure in children and adolescents. Nat Neurosci 18, 773– 778. 10.1038/nn.3983

Noble, K.G., Houston, S.M., Kan, E., Sowell, E.R., 2012. Neural correlates of socioeconomic status in the developing human brain. Dev Sci 15, 516–527. 10.1111/j.1467-7687.2012.01147.x

Nusslock, R., Brody, G.H., Armstrong, C.C., Carroll, A.L., Sweet, L.H., Yu, T., Barton, A.W., Hallowell, E.S., Chen, E., Higgins, J.P., Parrish, T.B., Wang, L., Miller, G.E., 2019. Higher Peripheral Inflammatory Signaling Associated With Lower Resting-State Functional Brain Connectivity in Emotion Regulation and Central Executive Networks. Biol Psychiatry 86, 153–162. 10.1016/j.biopsych.2019.03.968

Nusslock, R., Miller, G.E., 2016. Early-life adversity and physical and emotional health across the lifespan: A neuroimmune network hypothesis. Biol Psychiatry. 10.1016/j.biopsych.2015.05.017

Olson, L., Chen, B., Fishman, I., 2021. Neural correlates of socioeconomic status in early childhood: a systematic review of the literature. Child Neuropsychology 27, 390–423. 10.1080/09297049.2021.1879766

Orhun, G., Esen, F., Özcan, P.E., Sencer, S., Bilgiç, B., Ulusoy, C., Noyan, H., Küçükerden, M., Ali, A., Barburoğlu, M., Tüzün, E., 2018. Neuroimaging Findings in Sepsis-Induced Brain Dysfunction: Association with Clinical and Laboratory Findings. Neurocrit Care 30, 106–117. 10.1007/s12028-018-0581-1

Orhun, G., Tüzün, E., Bilgiç, B., Ergin Özcan, P., Sencer, S., Barburoğlu, M., Esen, F., 2020. Brain Volume Changes in Patients with Acute Brain Dysfunction Due to Sepsis. Neurocrit Care 32, 459–468. 10.1007/s12028-019-00759-8

Pearson, T.A., Bazzarre, T.L., Daniels, S.R., Fair, J.M., Fortmann, S.P., Franklin, B.A., Goldstein, L.B., Hong, Y., Mensah, G.A., Sallis, J.F., Smith, S., Stone, N.J., Taubert, K.A., 2003. American Heart Association Guide for Improving Cardiovascular Health at the Community Level: A statement for public health practitioners, healthcare providers, and health policy makers from the American Heart Association expert panel on population and prevention science. Circulation. 10.1161/01.CIR.0000054482.38437.13

Popescu, B.F.G., Lennon, V.A., Parisi, J.E., Howe, C.L., Weigand, S.D., Cabrera-Gómez, J.A., Newell, K., Mandler, R.N., Pittock, S.J., Weinshenker, B.G., Lucchinetti, C.F., 2011. Neuromyelitis optica unique area postrema lesions: Nausea, vomiting, and pathogenic implications. Neurology 76, 1229–1237. 10.1212/WNL.0b013e318214332c

Prinster, A., Quarantelli, M., Orefice, G., Lanzillo, R., Brunetti, A., Mollica, C., Salvatore, E., Morra, V.B., Coppola, G., Vacca, G., Alfano, B., Salvatore, M., 2006. Grey matter loss in relapsing-remitting multiple sclerosis: A voxel-based morphometry study. Neuroimage 29, 859–867. 10.1016/j.neuroimage.2005.08.034

Rao, U., Chen, L., Bidesi, A., Shad, M., Thomas, M., Hammen, C., 2010. Hippocampal changes associated with early-life adversity and vulnerability to depression. Biol Psychiatry 67, 357– 364.

Rondó, P.H., Pereira, J.A., Lemos, J.O., 2013. High sensitivity C-reactive protein concentrations, birthweight and cardiovascular risk markers in Brazilian children. Eur J Clin Nutr 67, 664–669. 10.1038/ejcn.2013.75

Rudolph, M.D., Graham, A.M., Feczko, E., Miranda-Dominguez, O., Rasmussen, J.M., Nardos, R., Entringer, S., Wadhwa, P.D., Buss, C., Fair, D.A., 2018. Maternal IL-6 during pregnancy can be estimated from newborn brain connectivity and predicts future working memory in offspring. Nat Neurosci 21, 765–772. 10.1038/s41593-018-0128-y

Rytter, M.J.H., Kolte, L., Briend, A., Friis, H., Christensen, V.B., 2014. The immune system in children with malnutrition - A systematic review. PLoS One 9. 10.1371/journal.pone.0105017

Sankowski, R., Mader, S., Valdes-Ferrer, S., 2015. Systemic inflammation and the brain: novel roles of genetic, molecular, and environmental cues as drivers of neurodegeneration. Front Neurosci 9, 1–20.

Schlenz, H., Intemann, T., Wolters, M., González-Gil, E.M., Nappo, A., Fraterman, A., Veidebaum, T., Molnar, D., Tornaritis, M., Sioen, I., Mårild, S., Iacoviello, L., Ahrens, W., 2014. C-reactive protein reference percentiles among pre-adolescent children in Europe based on the IDEFICS study population. Int J Obes 38, S26–S31. 10.1038/ijo.2014.132

Sharshar, T., Hopkinson, N.S., Orlikowski, D., Annane, D., 2005. Science review: The brain in sepsis - Culprit and victim. Crit Care 9, 37–44. 10.1186/cc2951

Sheridan, M.A., Fox, N.A., Zeanah, C.H., McLaughlin, K.A., Nelson, C.A., 2012. Variation in neural development as a result of exposure to institutionalization early in childhood. Proceedings of the National Academy of Sciences 109, 12927–12932. 10.1073/pnas.1200041109

Sheridan, M.A., McLaughlin, K.A., 2014. Dimensions of early experience and neural development: Deprivation and threat. Trends Cogn Sci 18, 580–585. 10.1016/j.tics.2014.09.001

Shimony, J.S., Smyser, C.D., Wideman, G., Alexopoulos, D., Hill, J., Harwell, J., Dierker, D., Van Essen, D.C., Inder, T.E., Neil, J.J., 2016. Comparison of cortical folding measures for evaluation of developing human brain. Neuroimage 125, 780–790. 10.1016/j.neuroimage.2015.11.001

Shuto, H., Kataoka, Y., Horikawa, T., Fujihara, N., &hi, R., 1997. Repeated interferon-a administration inhibits dopaminergic neural activity in the mouse brain, Brain Research.

Spann, M., Bansal, R., Hao, X., Rosen, T., Peterson, B., 2020. Prenatal socioeconomic status and social support are associated with neonatal brain morphology, toddler language and psychiatric symptoms. Child Neuropsychology 26, 170–188. 10.1080/09297049.2019.1648641.Prenatal

Sugimoto, K., Kakeda, S., Watanabe, K., Katsuki, A., Ueda, I., Igata, N., Igata, R., Abe, O., Yoshimura, R., Korogi, Y., 2018. Relationship between white matter integrity and serum in fl ammatory cytokine levels in drug-naive patients with major depressive disorder: diffusion tensor imaging study using tract-based spatial statistics. Transl Psychiatry 8, 1–8. 10.1038/s41398-018-0174-y

Tauman, R., Ivanenko, A., O’brien, L.M., Gozal, D., 2004. Plasma C-Reactive Protein Levels Among Children With Sleep-Disordered Breathing, PEDIATRICS.

Taylor, M.D., Allada, V., Moritz, M.L., Nowalk, A.J., Sindhi, R., Aneja, R.K., Torok, K., Morowitz, M.J., Michaels, M., Carcillo, J.A., 2020. Use of C-Reactive Protein and Ferritin Biomarkers in Daily Pediatric Practice Practice Gap. Pediatr Rev 41, 172–183.

Tiberio, M., Chard, D., Altmann, D., Davies, G., Griffin, C., Rashid, W., Sastre-Garriga, J., Thompson, A.J., Miller, D., 2005. Grey and white matter volume changes in early primary progressive multiple sclerosis: A longitudinal study. Brain 128, 1454–1460. 10.1093/brain/awh498

Tooley, U.A., Bassett, D.S., Mackey, A.P., 2021. Environmental influences on the pace of brain development. Nat Rev Neurosci 1–13. 10.1038/s41583-021-00457-5

Tottenham, N., Hare, T.A., Quinn, B.T., McCarry, T.W., Nurse, M., Gilhooly, T., Millner, A., Galvan, A., Davidson, M.C., Eigsti, I.M., Thomas, K.M., Freed, P.J., Booma, E.S., Gunnar, M.R., Altemus, M., Aronson, J., Casey, B.J., 2010. Prolonged institutional rearing is associated with atypically large amygdala volume and difficulties in emotion regulation. Dev Sci 13, 46–61. 10.1111/j.1467-7687.2009.00852.x

Turesky, T., Jensen, S., Yu, X., Kumar, S., Wang, Y., Sliva, D., Gagoski, B., Sanfilippo, J., Zöllei, L., Boyd, E., Haque, R., Kakon, S., Islam, N., Petri, W., Nelson, C., N, G., 2019. The relationship between biological and psychosocial risk factors and resting-state functional connectivity in 2-month-old Bangladeshi infants: a feasibility and pilot study. Dev Sci 22, e12841.

Turesky, T., Shama, T., Kakon, S.H., Haque, R., Islam, N., Someshwar, A., Gagoski, B., Petri, W.A., Nelson, C.A., Gaab, N., 2021. Brain Morphometry and Diminished Physical Growth in Bangladeshi Children Growing up in Extreme Poverty: a Longitudinal Study. Dev Cogn Neurosci 52, 101029. 10.1016/j.dcn.2021.101029

Turesky, T., Xie, W., Kumar, S., Sliva, D.D., Gagoski, B., Vaughn, J., Lilla, Z., Petri, W.A., Nelson, C.A., Gaab, N., 2020. Relating anthropometric indicators to brain structure in 2-month-old Bangladeshi infants growing up in poverty: A pilot study. Neuroimage 210, 1–10. 10.1016/j.neuroimage.2020.116540

Van Essen, D.C., 2005. A Population-Average, Landmark- and Surface-based (PALS) atlas of human cerebral cortex. Neuroimage 28, 635–662. 10.1016/j.neuroimage.2005.06.058

VanTieghem, M., Korom, M., Flannery, J., Choy, T., Caldera, C., Humphreys, K.L., Gabard-Durnam, L., Goff, B., Gee, D.G., Telzer, E.H., Shapiro, M., Louie, J.Y., Fareri, D.S., Bolger, N., Tottenham, N., 2021. Longitudinal changes in amygdala, hippocampus and cortisol development following early caregiving adversity. Dev Cogn Neurosci 48, 100916. 10.1016/j.dcn.2021.100916

Varatharaj, A., Galea, I., 2017. The blood-brain barrier in systemic inflammation. Brain Behav Immun 60, 1–12. 10.1016/j.bbi.2016.03.010

Vercellino, M., Plano, F., Votta, B., Mutani, R., Giordana, M., Cavalla, P., 2005. Grey matter pathology in multiple sclerosis. J. Neuropathology Experimental Neurology 64, 1101–1107. 10.1007/978-88-470-2127-3_9

Wen, D.J., Poh, J.S., Ni, S.N., Chong, Y.S., Chen, H., Kwek, K., Shek, L.P., Gluckman, P.D., Fortier, M. V., Meaney, M.J., Qiu, A., 2017. Influences of prenatal and postnatal maternal depression on amygdala volume and microstructure in young children. Transl Psychiatry 7, e1103. 10.1038/tp.2017.74

Wersching, H., Duning, T., Lohmann, H., Mohammadi, S., Stehling, C., Fobker, M., Conty, M., Minnerup, J., Ringelstein, E.B., Berger, K., Deppe, M., Knecht, S., 2010. Serum C-reactive protein is linked to cerebral microstructural integrity and cognitive function. Neurology 74, 1022–1029. 10.1212/WNL.0b013e3181d7b45b

Westhoff, D., Engelen-Lee, J.Y., Hoogland, I.C.M., Aronica, E.M.A., van Westerloo, D.J., van de Beek, D., van Gool, W.A., 2019. Systemic infection and microglia activation: A prospective postmortem study in sepsis patients. Immunity and Ageing 16, 1–10. 10.1186/s12979-019-0158-7

Xie, W., Jensen, S.K.G., Wade, M., Kumar, S., Westerlund, A., Kakon, S.H., Haque, R., Petri, W.A., Nelson, C.A., 2019a. Growth faltering is associated with altered brain functional connectivity and cognitive outcomes in urban Bangladeshi children exposed to early adversity. BMC Med 17, 1–11.

Xie, W., Kumar, S., Kakon, S.H., Haque, R., Petri, W.A., Nelson, C.A., 2019b. Chronic inflammation is associated with neural responses to faces in bangladeshi children. Neuroimage 202, 1–9. 10.1016/j.neuroimage.2019.116110

Yiu, G., He, Z., 2006. Glial inhibition of CNS axon regeneration. Nat Rev Neurosci 7, 617–627. 10.1038/nrn1956

Yoon, B.H., Jun, J.K., Romero, R., Park, K.H., Gomez, R., Choi, C.J.-H., I-O., K., 1997. Amniotic fluid inflammatory cytokines (interleukin-6, interleukin-1(beta), and tumor necrosis factor-(alpha)), neonatal brain white matter lesions, and cerebral palsy. Am J Obstet Gynecol 177, 19–26.

